# Alterations in topology, cost and dynamics of gamma-band EEG functional networks in a preclinical model of traumatic brain injury

**DOI:** 10.1101/2024.12.06.627187

**Authors:** Konstantinos Tsikonofilos, Michael Bruyns-Haylett, Hazel G. May, Cornelius K. Donat, Andriy S. Kozlov

## Abstract

Traumatic brain injury is a major cause of disability leading to multiple sequelae in cognitive, sensory, and physical domains, including post-traumatic epilepsy. Despite extensive research, our understanding of its impact on macroscopic brain circuitry remains incomplete. We analyzed electrophysiological functional connectomes in the gamma band using a preclinical model of blast-induced traumatic brain injury over multiple time points after injury. We revealed differences in small-world propensity and rich-club structure compared to age-matched controls, indicating functional reorganization following injury. We further investigated cost-efficiency trade-offs, propose a computationally efficient normalization procedure for quantifying cost of spatially embedded networks that controls for connectivity strength differences, and suggest metabolic drivers as a candidate for the observed differences. Furthermore, we employed a brain-wide computational model of seizure dynamics and attribute brain reorganization to a homeostatic mechanism of activity regulation with the potential unintended consequence of driving generalized seizures. Finally, we demonstrated post-injury hyperexcitability that manifests as an increase in sound-evoked response amplitudes at the cortical level. Our work characterizes for the first time gamma-band functional network reorganization in a model of brain injury and proposes potential causes of these changes, thus identifying targets for future therapeutic interventions.

## Introduction

Traumatic brain injury (TBI) affects millions annually (1) with post-injury phenotypes spanning sensory (2), physical (3, 4), cognitive (5, 6), and psychological (6) domains. Blast-induced traumatic brain injury (bTBI) is prevalent in war zones (7, 8) and has been coined the signature injury of warfare (9), affecting both military personnel and civilians (10, 11).

To date, a comprehensive understanding of how macroscopic TBI phenotypes emerge from the underlying pathophysiology remains elusive (12). At the microscopic scale, Cantu et al. (13) revealed signatures of an imbalance of excitation and inhibition (EI) as a precursor of post-traumatic epilepsy (PTE), a documented sequela of injury (14). Indeed, many studies have highlighted the vulnerability of inhibitory neurons to TBI (13, 15–19). Crucially, EI imbalance can lead to maladaptive plasticity as loss of inhibition and excessive firing can induce synaptic strengthening and if uncontrolled, this process can lead circuits into a pathological regime (20). Recent studies have begun to map the multiscale reorganization of cortical circuits (17, 20–22), while Frankowski et al. (21) identified a widespread restructuring of inhibitory networks following focal TBI.

Yet another consequence of EI imbalance and excessive neuronal firing following TBI is metabolic stress. More specifically, the restoration of altered ionic gradients necessitates the activity of sodium-potassium pumps, in turn leading to elevated consumption of adenosine triphosphate (ATP) to power this active process. This heightened demand for ATP, coupled with reported mitochondrial dysfunction (23, 24), can culminate in an energy crisis within neural tissue. Ultimately, this has the potential to alter the functional regime required for the brain to maintain a balance between energy cost and efficiency (25). This balance is typically associated with the so-called ‘small-world’ topology in brain networks, which is characterized by a combination of densely interconnected clusters of neighboring nodes (whereby the short distances incur low energetic costs) and the existence of short paths linking any two nodes in the network (called ‘geodesic distance’, and distinct from physical distance), which ensures efficiency in information transfer (26). This topology is thought to emerge due to the metabolic demands constraining the structuring of brain networks, and is posited to strike a balance between efficient information transfer and minimal wiring cost (25). Thus, it is expected to be disrupted due to metabolic stress following a TBI insult and indeed, studies have shown a deviation from small-world topology in this context (27–29). A relatively unexplored avenue of investigation involves conceptualizing these functional brain networks as spatially embedded, taking *physical* rather than *geodesic* node distances into account and calculating network cost, with a common assumption being that strong or long-distance coupling incurs higher costs (30, 31). This approach allows for characterization of the cost-efficiency of network organization, aligning with theories of how features like smallworld organization might emerge in biological networks (25). Another critical feature of brain networks is the presence of a highly interconnected core of hub nodes, often referred to as the ‘rich-club’ structure, which supports computations and synchronization across the entire network (31, 32). Due to its strong coupling density, this functional backbone necessitates substantial metabolic resources, leading to elevated costs (31), and could also become disrupted in the context of a TBI-induced metabolic crisis. Crucially, insights from computational modeling suggest that this topology can influence the propensity of a network to generate seizures, while the contribution of individual nodes to this behavior is influenced by their centrality, that is, their hub-like characteristics (33). These network measures might therefore also be disrupted following TBI, in the context of EI imbalance. It follows that despite progress in TBI research, powerful analytical tools such as network theory (34–38) and dynamic brain modelling (36, 39), as well as translationally relevant recording modalities with the potential to bridge species and spatial scales and link mechanisms and phenotypes of injury, have been grossly underappreciated in preclinical models. In this work, we endeavor to address this gap by characterising functional connectomes obtained in an earlier investigation using a translationally-relevant EEG recording technique within a rat model of bTBI (40). In that study, we observed aberrant functional connectivity patterns in the gamma frequency band (25-80 Hz) suggesting a potential reorganization of distributed networks. Here, we analyze the topology of these functional networks and introduce a novel metric for quantifying network cost that is invariant to differences in connectivity strength by means of normalization to equivalent null models. Key network organizational features such as small-world propensity (SWP) and rich-club structure were disrupted after bTBI. Network cost and its trade-off with efficiency showed a biphasic response with a low-cost configuration at 1 month post-injury followed by a high-cost one at 3 months post-injury, which was not discernible with the non-normalized metric and was associated with weight gain, suggesting a novel link to post-injury metabolism. We investigate the consequences of the altered network topology for whole brain dynamics through computational modeling and observe an increase in seizure risk at the later timepoint simultaneously driven by most nodes in the network. Finally, we provide evidence for hyperexcitability in our injury model reflected by increased auditory evoked potential amplitudes.

## Results

### Blast-Induced Changes in the Topology of Resting Functional Networks

Centrality measures showed an overall trend for decrease for the blast group (Figure 1). Betweenness centrality (number of shortest paths in a network that transverse that node) was significantly reduced at the global level after blast (Figure 1A, *p <* 0.05, main effect of group). Eigenvector centrality (the centrality of a node as a function of the centrality of its neighbors) showed a trend towards a similar effect of reduction for the injury group (Figure 1B) but did not reach significance. Similarly, betweenness centrality normalized to that of equivalent random networks, was not different between groups (Figure S1). No timepoint or interaction effects were detected for these network measures. In terms of whole-network measures, an increasing trend in small-world propensity was observed for the blast group (Figure 2A). The small-world topology combines the high clustering coefficients of regular and the short path lengths of random networks (26, 41). Hence we dissected small-world propensity into its constituents by analyzing the Δ_*C*_ and Δ_*L*_ components (see Materials and Methods) and revealed a reduced deviation from a regular network in terms of clustering for the blast group at 1 month post-injury. This was accompanied by a larger deviation from the path length of an equivalent random network compared to the other experimental groups (Figure 2B, C). This trend was confirmed for Δ_*C*_ by statistical testing (*p <* 0.05, main effect of injury group) but did not reach significance for Δ_*L*_. No timepoint or interaction effects were observed in any of these 3 metrics.

**Fig. 1.**
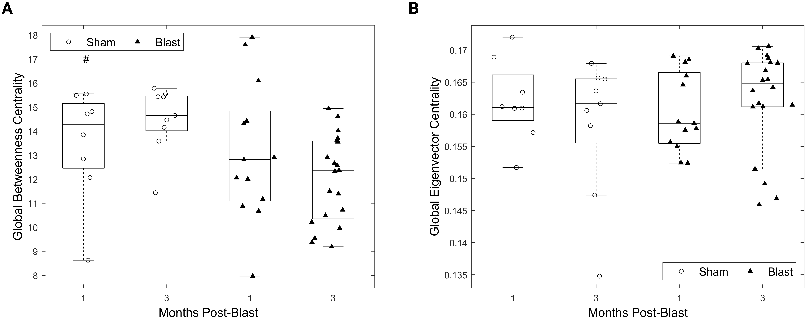
Decreased global centrality for the gamma connectivity networks chronically after injury. **(A)** Global betweenness centrality for all groups. **(B)** Global eigenvector centrality for all groups. **All panels:** Permutation-based 2-way ANOVA (Group x Time Post-Blast with interaction, 10000 permutations). Symbols denote *p*-values for main effect of group. #: *p <* 0.05 (blast 1-month: n = 13, blast 3-month: n = 20, sham 1-month n = 8, sham 3 -month n = 9).

**Fig. 2.**
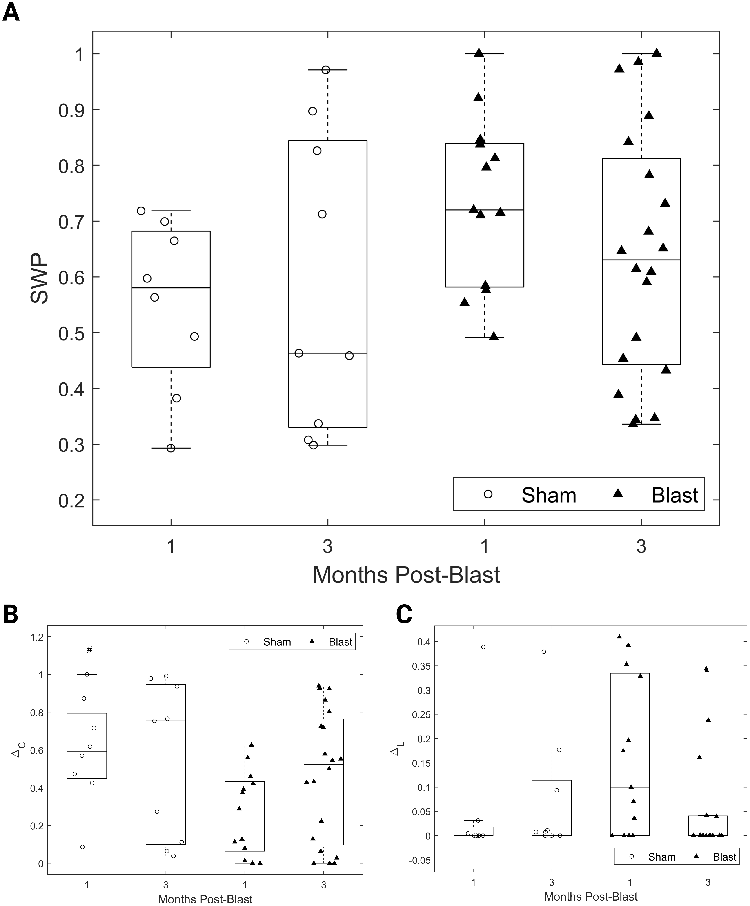
Decreased deviation from regular network clustering for the gamma connectivity networks after injury. **(A)** Small-World Propensity for all groups. **(B)** Fractional deviation from the mean clustering coefficient of an equivalent regular network for all groups. **(C)** Fractional deviation from the mean path length of an equivalent random network for all groups. **All panels:** Permutation-based 2-way ANOVA (Group x Time Post-Blast with interaction, 10000 permutations). Symbols denote *p*-values for main effect of group. #: *p <* 0.05 (blast 1-month: n = 13, blast 3-month: n = 20, sham 1-month n = 8, sham 3-month n = 9).

Finally, the presence of a rich-club structure was also assessed using the normalized rich-club coefficient (Figure 3). Over a range of richness parameter values a significant main effect of injury group was found with the blast groups having diminished rich-club structure compared to the sham groups.

**Fig. 3.**
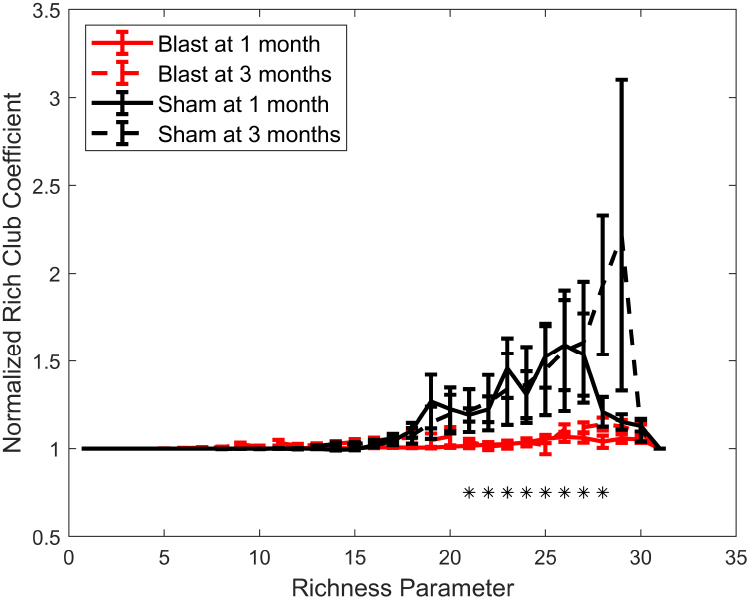
Loss of rich-club structure for the gamma connectivity networks chronically after injury. Permutation-based 2-way ANOVA (Group x Time Post-Blast with interaction, 10000 permutations) with FDR correction for multiple comparisons (n = 31 values of the richness parameter). Symbols denote corrected *p*-values for main effect of group. *\**: *p <* 0.05 (blast 1-month: n = 13, blast 3-month: n = 20, sham 1-month n = 8, sham 3-month n = 9).

### Blast-Induced Changes in Network Cost

The small-world topology is posited to achieve a favorable trade-off between cost and efficiency (25), while the rich-club structure is thought to incur high costs to a network (31). Hence, in an effort to identify putative drivers of the observed group differences in these topological features, we turn to the quantification of network cost. Network cost was first calculated similarly to Roy et al. (30) as the inner product of euclidean distances and connectivity values in the gamma band across node pairs. However, this cost metric could be spuriously driven by overall differences in connectivity between groups. Hence, we propose a novel normalization procedure based on equivalent null models (see Materials and Methods). Briefly, it consists of calculating lower and upper bounds of network cost for two inputs: a set of euclidean distances and a set of connectivity values. The normalized cost is then defined as the fractional deviation of the empirically observed cost from the minimum cost achievable given the inputs. This measure is robust to differences in connectivity strength and can be interpreted as reflecting a ‘strategy’ of resource (here connectivity strengths) allocation to a system of pre-specified properties (here node distances), a potential plasticity-based phenomenon at play in the post-TBI brain under metabolic distress. It is also computationally tractable since it only involves a sorting operation of the distance and connectivity sets and 3 inner product calculations.

Network cost was found to be elevated for the blast groups (*p <* 0.01, main effect of group, Figure 4A) when using the non-normalized cost metric of Roy et al. (30) for the gamma band. However, a different pattern of results emerged when considering our normalized cost measure for which we observed a significant interaction effect (*p <* 0.05, Figure 4B). Post-hoc pairwise comparisons revealed a significantly *decreased* normalized cost for the blast group at 1 month compared to the sham group at the same timepoint (*p <* 0.001, permutation-based *t*-test), and a significantly *elevated* normalized cost for the blast group at 3 months compared to the blast group at 1 month (*p <* 0.001, permutation-based *t*-test). The blast group at 3 months was not found to be significantly different to the sham group at 3 months.

**Fig. 4.**
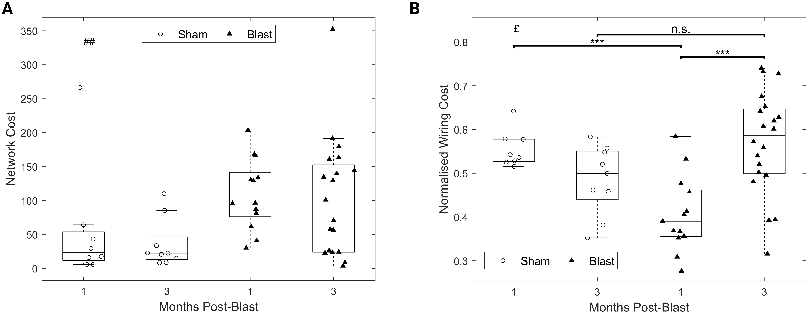
Dynamic changes in cost of gamma functional networks chronically after injury. **(A)** Non-normalized network cost, similar to Roy et al. (30). **(B)** Normalized network cost introduced here. **All panels:** Permutation-based 2-way ANOVA (Group x Time Post-Blast with interaction, 10000 permutations). Symbols denote *p*-values for main effect of group (#), interaction (£), or two-sample *t* -test (*\**). £: *p <* 0.05, ##: *p <* 0.01, *\* * \**: *p <* 0.001 (blast 1-month: n = 13, blast 3-month: n = 20, sham 1-month n = 8, sham 3-month n = 9).

Interestingly, our normalized cost metric was found to be significantly negatively correlated with normalized weight gain for the pooled blast groups (Figure 5, red hue symbols, Spearman *ρ* = − 0.52, *p <* 0.01), while being positively correlated for the pooled sham groups (Figure 5, blue hue symbols, Spearman *ρ* = 0.71, *p <* 0.01). The effect direction was preserved when correlations were examined in all 4 subgroups in isolation except for the blast group measured at 1 month (see Figure S2).

**Fig. 5.**
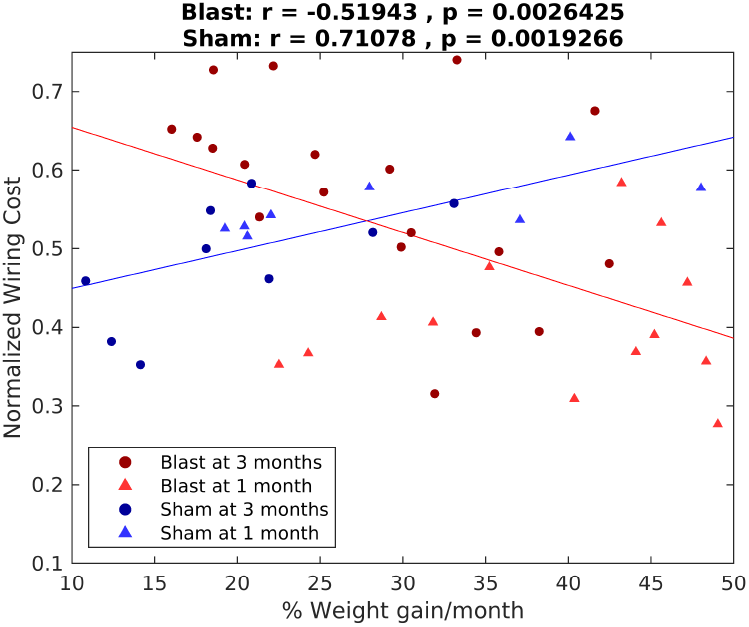
Opposite correlation signs between normalized network cost and normalized weight gain between sham and injury groups in the chronic phase. Spearman correlation values and associated *p*-values are reported. Lines correspond to best fit lines using least squares (red line: blast groups, blue line: sham groups), blast 1-month: n = 13, blast 3-month: n = 20, sham 1-month n = 8, sham 3-month n = 9.

These alterations in network cost following TBI could be a sign of broader changes in cost-efficiency trade-offs (25) in the injured brain. Hence, we further integrate our proposed cost measure in cost-efficiency metrics, using the fractional deviations of Small-World Propensity (see Materials and Methods) as proxies of network deviation from efficiency. Specifically, we interpret clustering deviation from the equivalent regular network as a measure of inefficiency of segregation, and we interpret path length deviation from the equivalent random network as a measure of inefficiency of integration. We then interpret high cost-efficiency as having low cost and efficiency deviations from their optimal values (see Materials and Methods). By virtue of their constituent quantities being normalized with respect to appropriate null models, these measures are also robust to differences in network density and overall connectivity strength.

A significant interaction effect was found for both segregative (Figure 6A, *p <* 0.05) and integrative (Figure 6B, *p <* 0.01) cost-efficiency and further post-hoc pairwise comparisons revealed increased cost-efficiencies for the blast group at 1 month compared to the sham group at the same timepoint (segregative: *p <* 0.001, Figure 6A, integrative: *p <* 0.05, Figure 6B), as well as compared to the blast group at 3 months (segregative: *p <* 0.01, Figure 6A, integrative: *p <* 0.01 Figure 6B). Finally, an effect of decreased cost-efficiency for the blast group at 3 months, compared to the sham group at the same timepoint was found for integrative cost efficiency (*p <* 0.05, Figure 6B).

**Fig. 6.**
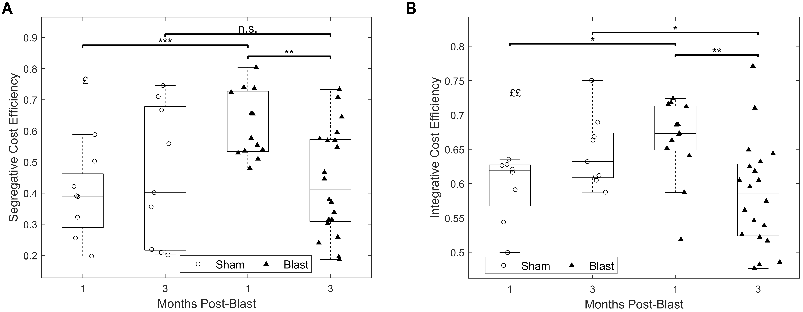
Dynamic changes in cost-efficiency of gamma functional networks chronically after injury. **(A)** Segragative cost-efficiency. **(B)** Integrative costefficiency. **All panels:** Permutation-based 2-way ANOVA (Group x Time Post-Blast with interaction, 10000 permutations). Symbols denote *p*-values for interaction effect (£), or two-sample *t* -test (*\**). £, *\**: *p <* 0.05, ££, *\*\**: *p <* 0.01, *\* * \**: *p <* .(blast 1-month: n = 13, blast 3-month: n = 20, sham 1-month n = 8, sham 3-month n = 9).

### Blast-Induced Changes in Ictogenicity and the Influence of Topology

To decipher the role of altered network topology on seizure dynamics, we employed the brain ictogenicity framework (33, 42, 43). Briefly, functional connectivity values inform the coupling strengths for a network model, wherein the node dynamics are governed by a phase oscillator model operating in either a resting or a rotating dynamical regime and can therefore be interpreted as resting or seizure dynamics, respectively (see Materials and Methods). Overall, stronger coupling would result in higher ictogenicity and vice versa (e.g., see Figure 4a in (33)). A control parameter tunes the excitability of individual nodes, and the whole network activity is simulated whilst parametrically varying this value. Outcome measures include the seizure probability (*P*_*sz*_), brain network ictogenicity (BNI), and node ictogenicity (NI), the latter quantifying the contribution of each node to seizure probability in the whole network.

Globally, an interaction effect was found for BNI (Figure 7A, *p <* 0.05) and post-hoc pairwise comparisons revealed an increased BNI for the blast group at 3 months postinjury compared to the corresponding sham group (Figure 7A, *p <* 0.05). Plotting seizure probability (*P*_*sz*_) over excitability (*I*_0_) values revealed that the increased BNI for the blast group at 3 months was owed to increased seizure probability at intermediate-to-high values of the excitability parameter (Figure 7B).

**Fig. 7.**
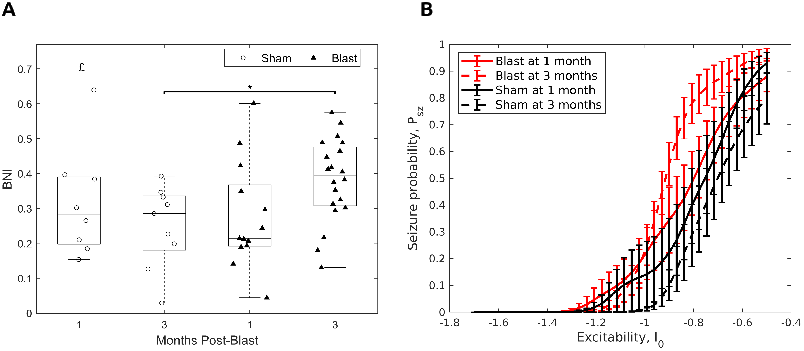
Increased network ictogenicity and seizure probability for the gamma functional networks 3 months post-injury. **(A)** Brain Network Ictogenicity. Permutation-based 2-way ANOVA (Group x Time Post-Blast with interaction, 10000 permutations). Symbols denote *p*-values for interaction effect (£), or two-sample *t* -test (*\**). £, *\**: *p <* 0.05 (blast 1-month: n = 13, blast 3-month: n = 20, sham 1-month n = 8, sham 3-month n = 9). **(B)** Seizure probability (*P*_*sz*_) as a function of excitability (*I*_0_).

NI quantifies the contribution of individual nodes to ictogenicity (Figure 8A). A significant effect of injury group was revealed for mean NI over the electrode array with decreased mean NI for the blast groups compared to shams (Figure 8B, main effect of group *p <* 0.05). For the standard deviation of NI over the electrode array a trend pointed towards a reduction for the blast groups across timepoints (Figure 8C).

**Fig. 8.**
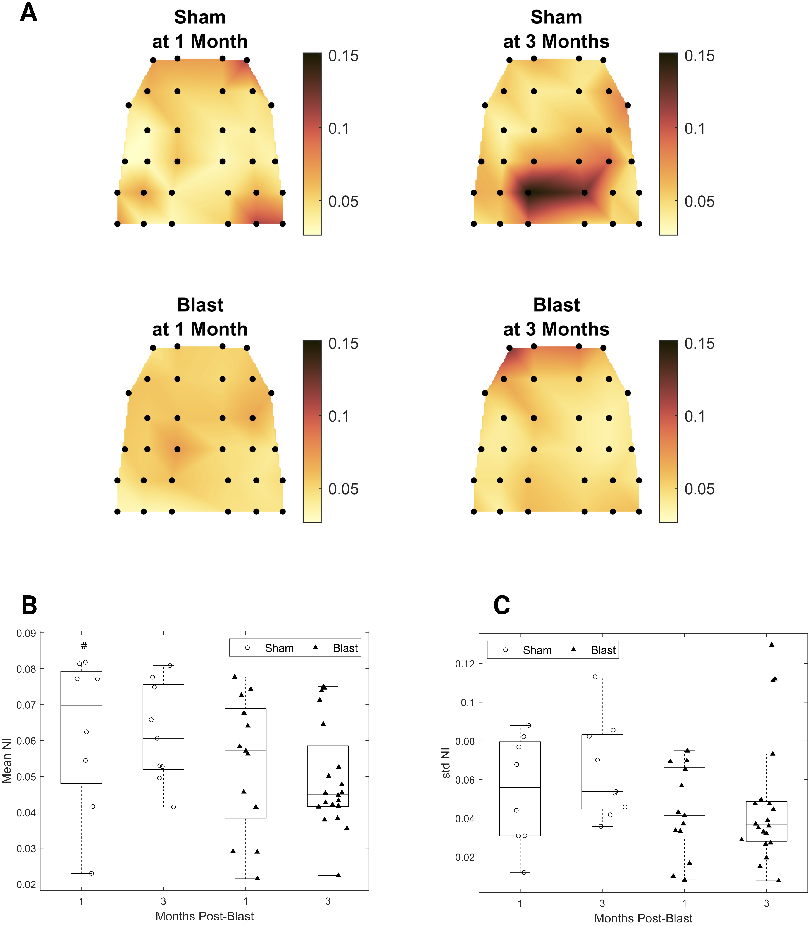
Decreased node ictogenicity for the gamma band functional networks chronically after injury. **(A)** Topoplots depicting mean group node ictogenicity. **(B)** Mean node ictogenicity. **(C)** Standard deviation of node ictogenicity. **(B, C):** Permutation-based 2-way ANOVA (Group x Time Post-Blast with interaction, 10000 permutations). Symbols denote *p*-values for the main effect of group. #: *p <* 0.05 (blast 1-month: n = 13, blast 3-month: n = 20, sham 1-month n = 8, sham 3-month n = 9).

As expected by the hub-like characteristics of a node to be a proxy for its influence on the dynamics of the whole network (33), NI of individual nodes strongly correlated with centrality measures (Figure S3A, Spearman *ρ* = 0.6, *p <* 0.001); Figure S3B, Spearman *ρ* = 0.85, *p <* 0.001). Furthermore, mean NI was correlated with the maximum value of the richclub coefficient across richness parameter values (Figure S4, *p <* 0.001, Adjusted R-Squared: 0.611).

### Blast-Induced Changes in Auditory Evoked Potentials

As supporting evidence for the presence of a disrupted EI balance in our injury model, a putative mechanistic basis underpinning the observed differences in topology and dynamics, we probed the excitability of central auditory regions using evoked potentials. For broadband clicks, grand averages of evoked potential waveforms indicated increased amplitudes for the blast group and were particularly pronounced at the 1 month timepoint (Figure 9). Analysis of the first negative component (N1) amplitude showed a significant interaction effect (*p <* 0.001, Figure 10), for which post-hoc pairwise comparisons revealed a significant increase of magnitude for the blast group at 1 month both compared to the sham group at the same timepoint (*p <* 0.01) and to the blast group at 3 months (*p <* 0.01). Similarly, increased amplitudes for the blast group were found for most other stimuli employed (Figures S5-S7).

**Fig. 9.**
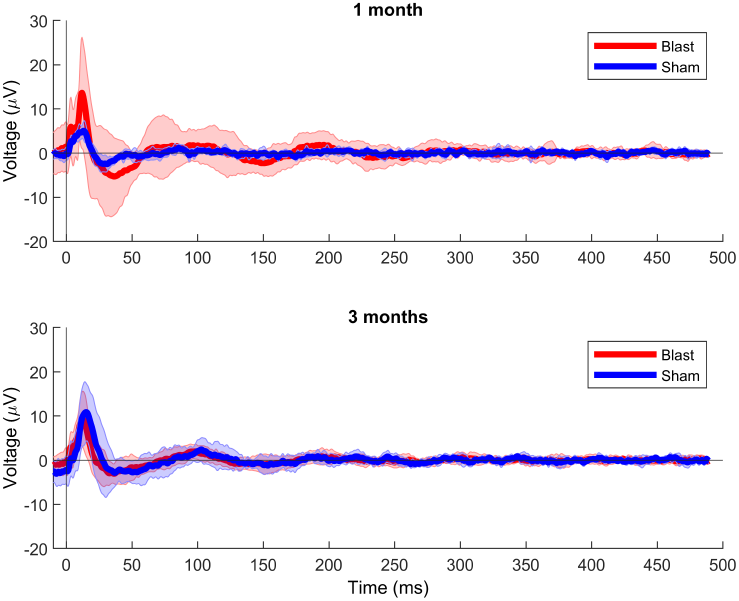
AEP waveforms for all groups - broadband clicks. Grand average waveforms for the 1-month (top) and 3-month groups (bottom). Shaded areas denote the standard deviation across the group. blast 1-month: n = 14, blast 3-month: n = 20, sham 1-month: n = 8, sham 3-month: n = 10.

**Fig. 10.**
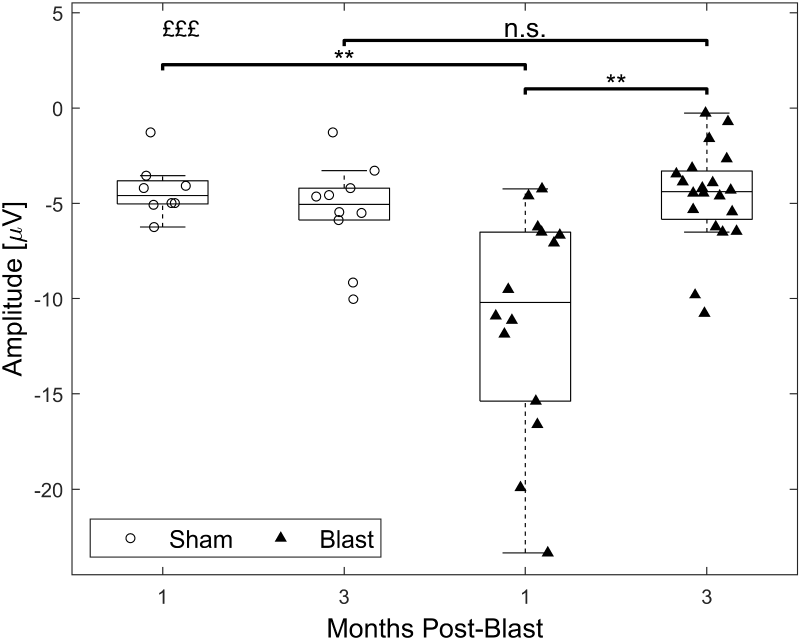
Increase in AEP N1 magnitudes 1 month after blast in response to broadband clicks. Permutation-based 2-way ANOVA (Group x Time Post-Blast with interaction, 10000 permutations). Symbols denote *p*-values for an interaction effect (£), or two-sample *t* -test (*\**). *\*\**: *p <* 0.01, £££: *p <* 0.001 blast 1-month: n = 14, blast 3-month: n = 20, sham 1-month: n = 8, sham 3-month: n = 10.

## DISCUSSION

In this study, we observed changes in the topology of gammaband functional networks after injury. At the local level, we found a decrease in betweenness centrality. At the global scale, networks exhibited an increase in small-world propensity, driven by increased clustering. Additionally, there was a pronounced loss of rich-club structure. We identified a downward trajectory of network costs (increased cost-efficiencies) at 1 month post-injury, which recovered by the third month post-injury, a pattern only discernible using our newly proposed, normalized metric. Importantly, network cost was associated with post-injury weight gain, with the correlation sign differing between groups: positive for the sham and negative for the blast cohorts. Furthermore, through computational modeling, we found heightened network ictogenicity at 3 months post-injury and a diminished role of individual nodes in determining overall epileptogenesis, a factor that showed significant correlation with graph-theoretical attributes such as centrality and presence of a rich-club structure. Finally, we provide evidence for post-injury hyperexcitability through increased amplitudes of auditory evoked potentials.

Our findings align with prior research. EEG-based studies employing these analytic tools in the context of TBI are scarce, yet altered small-worldness has been reported (29). In the MRI-based literature, increased path lengths have been observed after TBI (27, 28), echoing the trends observed in the present study. Notably, the localization of the patterns observed here within the gamma frequency range hints at a plastic reorganization of inhibitory networks (17, 22). Inhibitory neurons are known to shape activity in this band (44), and are impacted by injury in our model (40). Adding weight to this interpretation, Frankowski et al. (21) documented a similar trend in an animal model of focal non-blast TBI. Their findings illustrated increased local and reduced long-range input to inhibitory interneurons, even in areas distant from the injury locus. Strikingly, we observed this effect along with a disruption in the rich-club structure, in line with a diffusion imaging study (45). To the best of our knowledge, our work constitutes the first EEG study to demonstrate such a phenomenon within the scope of animal TBI research, while MEG studies in humans have demonstrated hemisphere and area dependent changes of rich-club hub representation (46) and hyperactivation of rich-club networks (47), with the latter result echoing observations in fMRI studies (48, 49).

To identify drivers of altered small-worldness, we probed a mechanism related to metabolism by characterizing the cost-efficiency trade-off thought to be afforded by the small-world topology. Several studies have aimed to quantify this tradeoff in terms of cost metrics that incorporate factors such as connection distance and strength (30, 31). Crucially, a recent modelling study by Achterberg et al. (50) suggests that artificial neural networks incorporating performance and cost terms in their objective functions to be optimised, converge to a small-world architecture. However, recent literature has emphasized the importance of using appropriate null models when studying network topology, as confounding factors such as weight distribution may spuriously drive group differences in topological measures (51)–a particularly relevant concern in the context of TBI-induced differences in connectivity strength. We sought to integrate these two lines of argument by introducing a novel, computationally-efficient normalization procedure for calculating cost in weighted networks based on equivalent null models. This has revealed a complex trajectory of post-TBI network cost in our model, which was not discernible using the standard metric. The trajectory of diminished cost in early chronic stages followed by an uptick in later stages might suggest a temporally evolving metabolic perturbation post-injury. A substantial body of literature connects TBI with metabolic aberrations (24, 52–57), noting a multiphasic response typified by an initial hypermetabolism phase succeeded by a subsequent hypometabolic phase and potential recovery to baseline levels (58). Importantly, post-TBI cerebral glucose metabolism as measured by positron emission tomography (PET) has been found to be positively correlated with full-scale Intelligence Quotient (fIQ) scores (59), further corroborating the existence of tradeoffs between metabolism and cognition.

Interestingly, there was a reversal in the direction of correlation between our proposed network cost metric and weight gain across groups. Since our metric can be interpreted as a proxy measure of resource allocation strategy (lower cost reflecting an allocation of resources that lies at a lower part of the cost range of possible allocations given the distances and connectivity strengths in the network), one explanation for this sign reversal hinges on potential differences of energy intake across cohorts due to the mediating effect of TBI. In a typical brain, increased energy supply might lay the foundation for allowing elevated network costs (i.e., a more ample allocation of resources), presumably promoting enhanced computational capabilities. In contrast, TBI-induced metabolic challenges might simultaneously and homeostatically drive heightened food intake and, in addition, employ a more frugal resource allocation strategy, depressing network costs. This would, in turn, render weight gain inversely associated with network cost, as demonstrated here. Likewise, the restored costs at 3 months post-injury could reflect a compensatory response. Specifically, the reduced cost at the 1-month mark might curtail computational capabilities enough to disrupt the organism’s normal function, compelling a shift to a more resource-intensive configuration to offset these processing challenges. Significantly, these findings on cost dynamics align with insights derived from cost-efficiency analyses, further buttressing this line of interpretation.

The localization of these results in the gamma frequency band is also relevant from a metabolic perspective, as activity of fast-spiking inhibitory interneurons has been shown to be energetically demanding (60). This has led to a hypothesis put forward by Kann (61), that these neural subtypes might be preferentially affected by TBI through an insult to metabolic processes. Our present and previous (40) findings are in line with this hypothesis and extend it to the macroscopic spatial scale and reorganization of brain-wide networks as measured by EEG electrodes.

The rich-club structure is postulated to function as a densely interconnected core that integrates information processed by peripheral components (31, 32). Hence, this functional backbone necessitates substantial metabolic resources at a high cost (31). Consequently, the metabolic narrative might also illuminate the observed post-injury disruption of this topological feature observed in this study. However, a complementary perspective emerges from our computational modeling, emphasizing dynamics over metabolism. Specifically, the rich-club’s capacity to facilitate activity propagation across the network can be a double-edged sword. While beneficial for information transfer in a healthy brain, this could, in the backdrop of TBI and its associated hyperexcitability, instigate uncontrolled activation cascades, possibly culminating in seizures. Indeed, Lopes et al. (33) demonstrated that targeted elimination of rich-club nodes curtails network ictogenicity *in silico* and this intervention also corresponded with favorable outcomes in post-resective surgery of epileptic patients. Thus, the loss of rich-club organization post-TBI observed here might signify an adaptive strategy to counteract excessive activity spread and mitigate seizure risks, especially amidst EI imbalances. Our findings indicate that network-wide seizure probability remains unchanged at one month but is altered by the third month post-injury. This could reflect the interplay of shared (like rich-club organization) and distinct (small-worldness or cost) topological attributes across the post-injury timeline. Finally, the switchlike behaviour observed in the later timepoint, combined with the decreased contribution of individual nodes to seizures across the whole network, could suggest a potential substrate for generalized as opposed to focal seizures as observed in a study using data from clinical populations with these two epilepsy types (42). Nevertheless, the reduction in individual node contribution as well as its correlation with topological measures lends support to the interpretation of an adaptive loss of rich-club structure. Intriguingly, a similar pattern in node-level contributions has been identified in research focusing on Alzheimer’s disease (43), suggesting a potentially general adaptive mechanism applicable across diverse neurological conditions.

The influence of injury on evoked potentials was prominent, pointing towards hyperexcitability (62) in cortical auditory regions and echoing observations of increased amplitudes in paediatric epileptic patients (63). While focusing on a specialized subnetwork, we propose that this pattern, combined with disruptions to inhibitory interneurons across multiple brain regions within the same model (40), reinforces the validity of our interpretation of the observed network differences as indicative of a disrupted EI balance (12). Even though we cannot rule out a hearing-loss related driver of this phenotype (64, 65), recent research in a model of Fragile X syndrome (66) also noted a similar trend of increased amplitudes of auditory evoked potentials. Given that this syndrome shares a mechanistic substrate of hyperexcitability, it is likely that a significant portion of the effects observed here stem from bTBI.

Overall the graph theoretical characterization of the functional networks has yielded a nuanced view of the consequences of injury and, when supplemented by a new methodological contribution in calculating network cost that shows promise of increased sensitivity to experimental effects while controlling for potential confounds, has integrated electrophysiology with the metabolic component of TBI for the first time. Furthermore, the results of computational modeling have suggested that such alterations might be the brain’s homeostatic response to maintain its dynamics post-injury. The study also reiterates the dynamic nature of injury progression and the potential role of the inhibitory plasticitome (67), in line with a recent hypothesis that TBI causes a reopening of the early developmental critical period where the EI balance has not yet reached its stable state and plastic changes occur at a higher frequency and intensity (68).

Future work could record activity from awake behaving rodents (66, 69, 70). However, there is evidence to support the robustness of gamma-band connectivity differences to anaesthetic state (71) as well as the similarity of fMRI-based functional connectivity patterns under the anaesthetic agent used here (urethane) to the ones observed during wakefulness (72). Moreover, the suspected metabolic disruptions in this model and their association with the proposed network cost metric should be validated by future studies using additional techniques able to interrogate brain metabolism such as PET scanning. Finally, by benchmarking this metric using larger datasets from clinical cohorts, its applicability can be gauged against other electrophysiological markers previously found to correlate with metabolism (73, 74).

## MATERIALS AND METHODS

The acquisition of the dataset analyzed here is described in our prior work (40), which provides detailed descriptions of the methodologies employed. All animal experiments were conducted in compliance with the Home Office license ([Scientific Procedures] Act 1986 and EU legislation). Briefly, Male Sprague Dawley rats were used, under a 2-by-2 factorial design based on injury group (sham or blast) and postinjury timepoint (1 or 3 months). Blast exposure was conducted using a **shock-tube**, inducing a mild-to-moderate injury. Electrophysiological recordings involved terminal anesthesia under urethane, and EEG recordings using Neuronexus electrodes and a SmartBox Pro/Radiens Allego system (Neuronexus, Michigan, USA). More details can be found in the Supplementary Materials. Follow-up analyses comprising this work are described in full detail below.

### Analysis of Functional Connectomes

#### Local Topological Measures

Graph theoretical analysis of resting-state functional connectivity networks (for the derivation of functional connectomes, see Supplementary Materials) was conducted using custom scripts, Matlab functions, functions from the Brain Connectivity Toolbox (BCT) (75) and publicly available code (41).

To characterize structure at the local level, node centrality measures were employed, quantifying the importance of a node in the network. *Betweenness centrality* (Matlab function *centrality*) is defined as the number of shortest paths in the network that transverse that node, while *eigenvector centrality* (BCT function *eigenvector_centrality_und*) measures the centrality (importance) of a node in a graph as a function of the centrality of its neighbors (76).

Global variants of node-specific measures were calculated by taking the mean across the electrode array. Since the magnitude of betweenness centrality can be influenced by additive shifts of the weight distribution, a normalized measure was also computed by dividing the global centrality of the observed networks over the mean global centrality of 100 equivalent random networks.

#### Global Topological Measures

To characterize topology at the *global* level, we first employed the Small-World Propensity (SWP) measure (41), which quantifies the small-world characteristics of a network by positioning it along the continuum from regular to random (as in the original formulation from Watts and Strogatz (26)) using appropriate null models (observed, equivalent regular, and equivalent random networks). More specifically, this measure quantifies the deviation of the clustering coefficient Δ_*C*_, from an equivalent regular network, and the deviation of path length Δ_*L*_, from an equivalent random network and is computed according to the following equation:

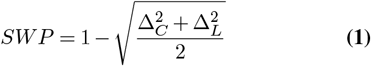

The deviations are calculated as follows:

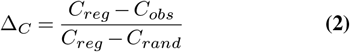

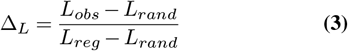

The subscripts *obs, reg, rand* refer to the observed, equivalent regular, and equivalent random networks, respectively. To calculate the clustering coefficient for individual nodes, the measure introduced by Onnela et al. (77) for weighted networks is used. For the calculation of path length, the weight of an edge is derived from the strength of functional coupling between the adjacent nodes, whereby nodes connected by a strong edge can be interpreted as being in close proximity from a neural communication perspective. Hence, following Muldoon et al. (41), the inverse of the edge weight is interpreted as distance. The mean minimum path length across node pairs and clustering coefficient across nodes are used to characterize the whole network. For more details on the calculations of network measures involved in small-world propensity, see Supplementary Materials.

The presence of rich-club structure was assessed using the approach by Van Den Heuvel et al. (78). Specifically, the rich-club coefficient was calculated for each value of the richness parameter defined as the node degree (BCT function *rich_club_wu*) and the same calculation was performed for 100 equivalent random networks (BCT function *randmio_und_connected*, which ensures that the randomized network maintains connectedness). Intuitively, rich-club structure can be understood as the presence of strong coupling between the hub (i.e. rich in connections) nodes of a network. Finally the *normalized* rich-club coefficient was calculated as the ratio of the observed coefficient to the mean of the coefficients of the random networks, for each value of the richness parameter.

#### A Novel Normalization Procedure for Cost Quantification

The concept of network cost has been particularly illuminating in the context of TBI. A notable study (30) determined network cost by computing the dot product of the functional connectivity matrix and the physical distance matrix, based on the premise that connections which are strongly coupled or spatially distant demand higher energetic expenditure. In the context of TBI, as with the network topology measures previously discussed, variations in cost—given a network’s spatial configuration (physical distances between nodes)—might trivially arise simply due to disparities in overall connectivity strength. Consequently, to draw valid comparisons between networks, it is necessary to employ a suitable normalization method or a null model. To this end, we introduce the concept of Total Effective Wiring Cost (TEWC). This metric is derived by initially determining the cost of the observed network, following the methodology of Roy et al. (30), where the cost is the dot product of edge weights and corresponding Euclidean distances between the adjacent nodes:

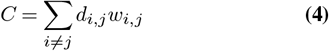

where *d*_*i,j*_ denoted the node distances and *w*_*i,j*_ the corresponding edge weights.

Following this, two surrogate null networks are constructed via reassignment of edge weights, representing the extremes of the cost spectrum for the given configuration of the network: the *minimum cost equivalent network* and the *maximum cost equivalent network*. Constructed to incur either the minimal or maximal cost, these networks maintain the sets of physical distances and connection strengths between nodes, and their costs can be calculated using Equation 4. Subsequently, the *total effective wiring cost* is defined as the fractional deviation from the minimum cost, relative to the maximum cost attainable:

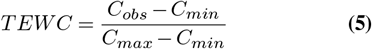

Where the subscripts *obs, min, max* refer to the observed, maximum cost equivalent, and minimum cost equivalent networks, respectively.

This formulation can be interpreted in terms of a *strategy* of resource allocation (i.e., weights) across a set of nodes with fixed physical distances. It is thus suitable to quantify potential network reorganization schemes that may be implemented after injury by plasticity mechanisms, for example as an adaptive response to a post-TBI energy crisis (24, 52– 56, 79, 80).

This reduction of the problem to one of edge allocation naturally leads to an efficient implementation through a closed form solution making use of the rearrangement inequality (81). Given two equally sized sets of real numbers *x*_1_, *x*_2_, .., *x*_*n*_ and *y*_1_, *y*_2_, …, *y*_*n*_ sorted in ascending order, the following inequality holds:

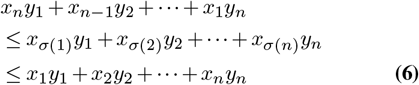

for every permutation *σ* of the indices 1, 2, …, *n*.

It follows that the cost calculation for the equivalent maximum and minimum cost networks described above reduces to sorting the weights of the observed network in descending and ascending order and taking their dot product with the sorted distance vector according to ascending order, yielding the cost of the minimum and maximum cost equivalent networks, respectively. Having a closed form, this metric is computationally efficient and reduces to one sorting operation per network. Our implementation makes use of the *sort* function of the core Matlab package, which in turn employs an implementation of the Quick Sort algorithm (82). This algorithm has an average performance of 𝒪(*n* log *n*), pointing to the scalability of TEWC for parcellations involving a large number of brain regions.

The brain’s organization does not strictly minimize cost (25), instead striking a balance between efficient organization and information processing capacity. Consequently, cost metrics are often evaluated within the framework of cost-efficiency (83), quantifying this aforementioned equilibrium.

Here, we introduce two metrics of cost efficiency by combining the newly introduced TEWC with the Δ_*C*_ and Δ_*L*_ deviations, yielding segregative and integrative cost-efficiency (SCE and ICE, respectively) as described below:

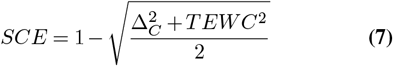

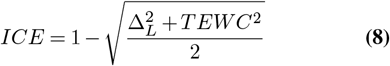

Analogous to how SWP gauges the coexistence of short path lengths with pronounced clustering, these cost-efficiency metrics evaluate the presence of low cost alongside computational capacity. Here, higher computational capacity is quantified as smaller deviation from the null equivalent models encompassed by the SWP framework, namely the equivalent regular and random networks. This in turn allows for a systematic decomposition of cost-efficiency into components associated with segregation and integration.

### Connectivity-Informed Whole Brain Computational Modeling

#### Simulations Using a Phase-Oscillator Model

Node dynamics were simulated using the following theta model (84):

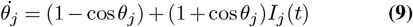

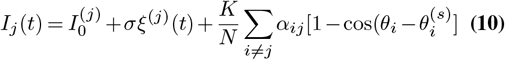

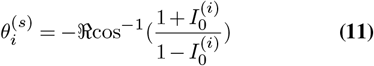

Where *θ*_*j*_ is the phase of node *j, I*_*j*_(*t*) the input current received by node *j* at time *t*, 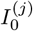 the excitability of node *j, ξ*^(*j*)^(*t*) noisy inputs to node *j* from distant regions, *α*_*ij*_ the coupling strength between nodes *i, j, K* a global scaling constant, *N* the number of nodes and 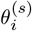 the stable phase of the oscillators in the absence of coupling and noise. Simulations were run for a total of 6 million timesteps and repeated 5 times, while the values of the excitability parameter *I*_0_ were varied.

Values for *K* and *I*_0_ were selected similarly to (43) and connectivity matrices were scaled to [0,1] by division with the maximum value. The specific parameter values used are detailed in Table 1.

**Table 1.** Parameters of the Computational Model of Brain Ictogenicity.

| Parameter | Meaning | Value |
| --- | --- | --- |
| $I_0$ | Node Excitability | $[-1.7, -0.5]$ |
| $K$ | Global Scaling of Coupling Strength | 10 |
| $A$ | Connectivity Matrix | dwPLI values scaled to $[0, 1]$ |
| $N$ | Network Size | 32 |
| $\sigma$ | Noise Level | 6 |
| $T$ | Simulation Steps | $4 \cdot 10^6$ |
| $\delta t$ | Simulation Time Step | $10^{-2}$ |

#### Outcome Measures of Ictogenicity

Following the simulations, two summary measures of the dynamics were calculated. Globally, BNI is defined as the integral of the seizure probability over the excitability parameter values (33):

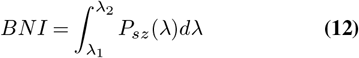

Where the seizure probability is defined as the proportion of simulation time during which the system is engaged in epileptic dynamics:

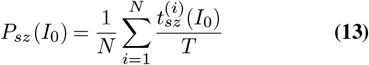

At the node level, NI quantifies the contribution of each node to the whole network’s BNI:

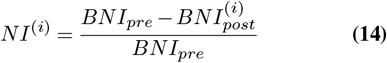

Where *BNI*_*pre*_ and *BNI*_*post*_ are the BNI of the full network and the BNI following the removal of node *i* respectively. Importantly, NI stands as a normalized metric, ensuring that potential group effects are not spuriously driven by differences in global connectivity strength.

#### Evoked Potentials Analysis

For the analysis of AEPs, the approach and terminology used by Jonak et al. (66) is adopted. For each electrode, the N1 component was automatically identified using the Matlab function *findpeaks* within a predetermined time window (15 to 100 ms post-stimulus). Subsequently, peak amplitude was extracted and stored for group-level analysis.

#### Statistical Analysis

Permutation-based 2-way ANOVA tests (85, 86) were conducted, including terms for the main effect of group, main effect of timepoint and an interaction effect. This procedure is robust to deviations from normality and outlier values, but it is not unaffected by violations of homoscedasticity. It can be argued, however, that variance changes in the outcome measures of an experimental group (particularly the injury group) is indeed an effect of interest and sensitivity to it is desirable.

In the case of a significant interaction effect, *post-hoc* pairwise comparisons were performed for the contrasts of interest (blast at 1 month vs blast at 3 months, blast at 1 month vs sham at 1 month, blast at 3 months vs sham at 3 months), using permutation-based two sample *t*-tests (Matlab function *ttest2* from the statistics and machine learning toolbox).

Correlations between continuous variables were assessed using Spearman correlation (Matlab function *corr*). In the presence of outlier observations, a robust regression approach was followed (Matlab function *fitlm* with default parameters). For tests examining group pairs for post-hoc pairwise analyses, correction for multiple comparisons was performed using the Bonferroni correction method (87). When considering values within a parameterized space (such as the richness or excitability parameters), the Benjamini-Hochberg FDR method was used (88). The threshold for significance was set at 0.05 for adjusted *p*-values in all tests.

In all figures, ‘#’ denotes a main effect of injury group, while ‘%’ denotes an interaction effect. ‘*’ denotes a significant effect in two-sample *t*-tests. The number of symbols has the usual interpretation (1 symbol: *p <* 0.05, 2 symbols: *p <* 0.01, 3 symbols: *p <* 0.001).

## ACKNOWLEDGEMENTS

This study was funded by the Royal British Legion at the Centre for Blast Injury Studies, Imperial College London, and by the US Army Research Office.

## Supplementary Note 1: Subjects and Experimental Design

Subjects were male Sprague Dawley rats, sourced from Charles River Laboratories, UK. Upon arrival at the animal holding facility, they were maintained under standardized conditions with *ad libitum* access to rodent chow and sterilized water and a regular 12-hour light/12-hour dark cycle. There were allocated into 4 subgroups according to a 2-by-2 between subjects factorial design, with injury group (sham or blast) and post-injury timepoint (1 or 3 months post injury) as the factors. All procedures were conducted in accordance to the Animals (Scientific Procedures) Act under appropriate licenses, and in consultation with named veterinary surgeons, as well as named animal care and welfare officers.

## Supplementary Note 2: Blast Injury

Blast injury was delivered utilizing the shock-tube located at the Centre for Blast Injury Studies at Imperial College London (89– 95). We have previously shown that this mild-to-moderate injury elicits hallmark electrophysiological and histopathological phenotypes (40). Anesthesia was achieved via isoflurane and physiological parameters were monitored over a 20-minute period until stabilized, where stability was defined as the absence of pedal reflexes, a respiratory rate within the range of 50-60 breaths per minute, and an oxygen saturation exceeding 95%. While under continuous isoflurane delivery, rats were positioned on a steel platform attached to the open end of the compressed air-driven shock-tube. Orientation was lateral, with the right side directed towards the incoming shock-wave. Pressure waveforms captured using piezoelectric sensors positioned at the shock tube’s exit, reflected a pattern consistent with the Friedlander model (96) with a peak overpressure of 232.5826(6.4486) kPa and positive phase duration of 1.3870(0.0427) ms (both values mean(sd)). Animals in the sham group underwent the same anesthetic protocol except for the blast exposure. Following the blast/sham procedure, animals were recovered in a heated cage and given peri-operative buprenorphine analgesia for 72 hours (pre-blast: 0.05 mg/kg via subcutaneous injection, post-blast: *ad libitum* self-administration of 0.3 mg/kg buprenorphine jelly). Full details of the blasting and peri-operative procedures can be found in (40).

## Supplementary Note 3: Electrophysiological Recordings

Terminal anaesthesia was induced using isoflurane, followed up by an introperitoneal injection of urethane (1.35 g/kg) for maintentance. Upon loss of the pedal reflex (usually within 1 hour), local anaesthetic was administered in the scalp area (bupicavaine 1.5 mg/kg) and atropine was administered subcutaneously (0.66 ml/kg, 1% w/v) to reduce mucous secretions. Subsequently, animals were transferred to a stereotaxic frame for surgery. A midline incision was made in the scalp using a scalpel, followed by blunt dissection of the connective tissues using scissors, clearing and smoothing of the dorsal surface of the skull with a blunt dental drill. All recordings were performed inside a shielded, anechoic chamber using Neuronexus electrodes (Rat EEG Functional), headstages (Neuronexus, SmartLink), and data acquisition system/software (SmartBox Pro/Radiens Allego). The EEG array was referenced to the nuchal musculature. Signals were bandpass filtered between 1.1 Hz and 15 kHz and sampled at 30 kHz.

## Supplementary Note 4: Resting-State Data Preprocessing

Data preprocessing was performed using functions from the EEGLAB Toolbox (97) and custom scripts in Matlab (version 2019b). Raw data were imported into the workspace, converted into EEGLAB format and downsampled to 1 kHz (function *pop_resample*) to reduce computational load. Subsequently, line noise was suppressed through the *CleanLine* plugin (98, 99) (function *pop_cleanlinenoise*) applied twofold to the 50 Hz line frequency and its harmonics. The two applications were performed using two different moving window sizes, which was empirically found to be successful in eliminating line noise from the acquired data. Data were also subjected to automatic artifact rejection (function *clean_artifacts*). Data were then visually inspected to confirm the success of this automated step. As a final preprocessing step, data were decomposed using the Infomax ICA algorithm (function *pop_runica*) to seperate artifactual components. Components were visually inspected in terms of their temporal activation and spectra and components containing primarily respiratory, cardiac or muscle-related artifacts were conservatively removed from the data. Datasets were excluded from further analysis in cases where line noise was not effectively suppressed, or the automated artifact rejection procedure led to more than 20% of the datapoints being rejected. 1 animal in the blast 1-month group and 2 animals in the sham 3-months group were excluded based on these criteria.

## Supplementary Note 5: Phase-Based Functional Connectivity

Functional connectivity was assessed via the debiased weighted phase-lag index (dwPLI) (100), a frequency-resolved, phase-based metric that quantifies the consistency of phase differences between two time series in a time-resolved manner, using Matlab and Fieldtrip functions. The weighting of each phase difference by its magnitude in the computation of dwPLI serves to eliminate zero-phase connectivity, which might be attributed to volume conduction (100), making it suitable for sensor-level connectivity analysis. The calculation involved segmenting the signals into windows of 3 s duration and then computing the cross-spectral density for each electrode pair using Welch’s method with a 1.5 s sliding window and 50% overlap (MATLAB function *cpsd*). These cross-spectra are subsequently used as input to the low-level the FieldTrip function *ft_connectivity_wpli*. Connectivity values for a frequency band are determined by averaging across the frequency bins specific to that band (gamma: 25-80 Hz).

## Supplementary Note 6: Small-world Propensity Calculation

To calculate the clustering coefficient for a node, the measure introduced by (77) for weighted networks is used:

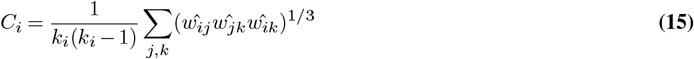

Where *C*_*i*_ is the clustering coefficient of node *i, k*_*i*_ the number of edges adjacent to node *i*, and 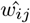 the weight of the edge between nodes *i* and *j* divided by the maximum weight of the network.

For the calculation of path lengths, the weight of an edge is derived from the strength of functional coupling between the adjacent nodes, hence nodes connected by a strong edge can be interpreted as being in close proximity from a neural communication perspective. Hence, following (41) the inverse of the edge weight is interpreted as distance. Shortest paths between any two nodes are calculated using Dijkstra’s algorithm and the mean minimum path length across pairs is used to characterise the whole network:

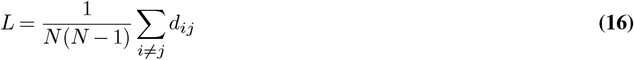

Where *N* is the number of nodes and *d*_*ij*_ the length of the shortest path connecting nodes *i* and *j*.

To generate an equivalent random network, edges from the observed network are randomly re-assigned among node pairs. For equivalent regular networks, edges from the observed network are organized in descending order and are then distributed along the subdiagonals of the adjacency matrix. These equivalent networks retain the weight distribution of the original network, making them appropriate normalization methods in the context of TBI.

## Supplementary Note 7: Weight Gain

In order to assess physiological correlates of the normalised network cost metric, its association with weight gain was explored. In an effort to account for the effect of longer post-injury survival of the 3-month post injury group the weight gain in g (percent of weight at the time of blast gained by the time of the electrophysiology experiment) was divided by the number of months post-injury, yielding a weight gain per month measure.

## Supplementary Note 8: Auditory Stimuli

Auditory stimuli were generated using Matlab at a sampling rate of 250 kHz and saved as *wav* files. Broadband clicks were created as rectangular pulses with a duration of 0.1 ms. Tone pips were generated as sinusoidal waves with a duration of 25 ms and a 5 ms cosine squared ramp. The tones had frequencies of 10, 37.5, and 65 kHz at a fixed intensity. All stimuli had an interstimulus interval of 500 ms and were presented 1000 times with alternating polarities to ensure artifact-free data with sufficient signal-to-noise ratio. All stimuli were delivered through dynamic speakers (Avisoft, Vifa) interfacing with Ultra-SoundGate Player 116 (Avisoft) for playback and digital to analog conversion at 16 bits, and software (Avisoft, RECORDER USGH) running on a Windows computer. Speakers were positioned at a distance of 22 cm from the animal’s ears. The intensity of the stimuli was calibrated to be 80 dB SPL at the ear level of the animal using an Avisoft CM24/CMPA condenser ultra-sound microphone, previously calibrated using a sinusoidal 40 Hz reference signal generator (Avisoft). The synchronization of sound stimuli and electrophysiological recordings was achieved through analog TTL pulses sent from Player 116 to the data acquisition module (Neuronexus SmartBox Pro).

## Supplementary Note 9: Auditory Stimulation Data Preprocessing

The preprocessing of data during auditory stimulation closely resembled that of resting-state data. Signals were downsampled to 2 kHz (function *pop_resample* with default settings, including an anti-aliasing filter at the new Nyquist frequency). Line noise was removed using the CleanLine plugin. An additional artifact rejection step involved identifying high-amplitude voltage excursions exceeding 400 *μV*, which were subsequently excluded. Subsequently, data were subjected to finite impulse response filtering (function *pop_eegfiltnew* with default parameters) between 3 and 300 Hz (cascade of low followed by highpass Hamming windowed sinc filters, orders: 88 and 3300 respectively). Finally data were segmented into epochs based on event markers (−10 to 490 ms, function *pop_epoch*) and the evoked potential waveforms extracted by taking the mean over trials.

## Supplementary Note 10: Supplementary Figures

**Fig. S1.**
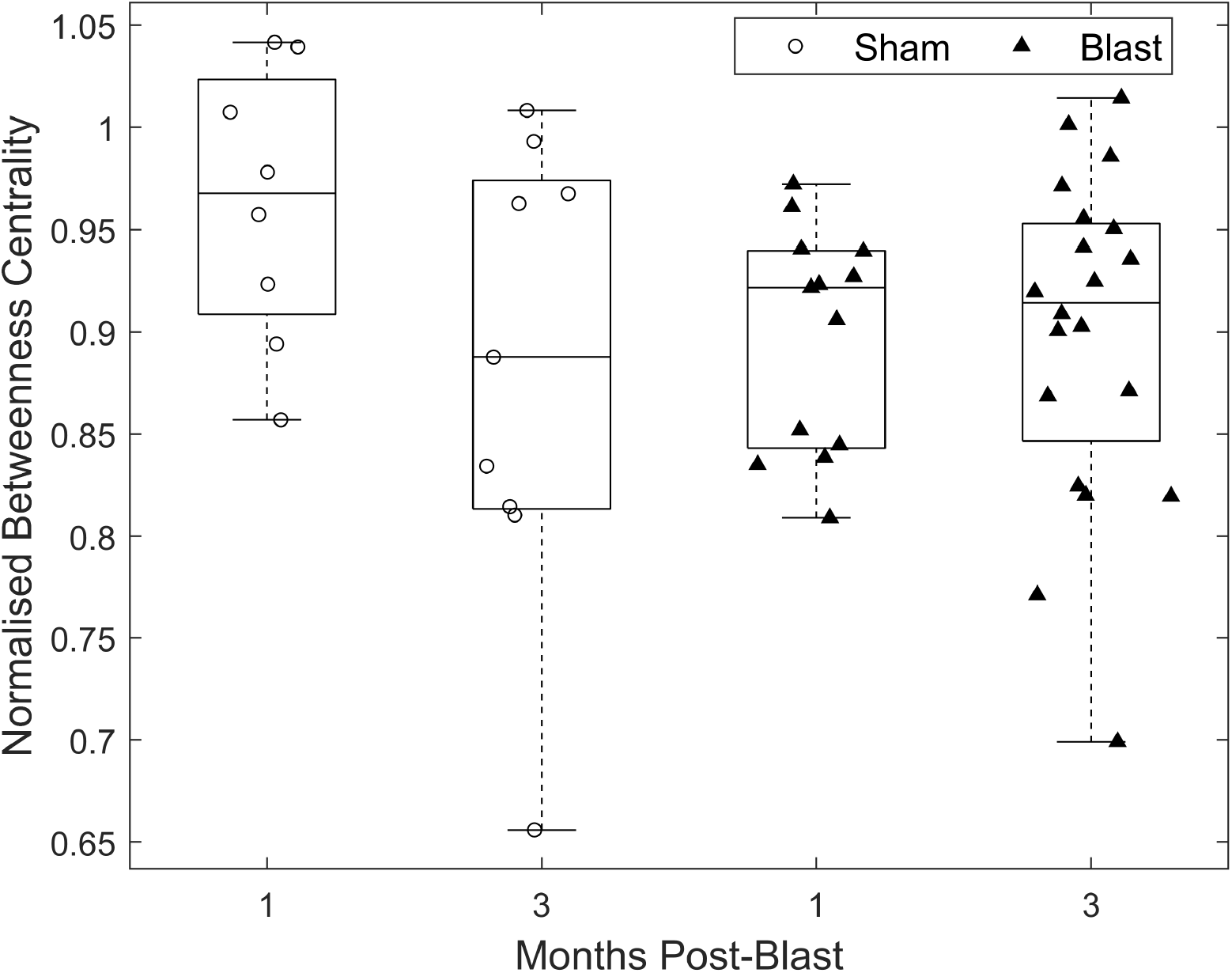
No effect on normalised global betweenness centrality of gamma functional networks chronically after injury. Permutation-based 2-way ANOVA (Group x Time Post-Blast with interaction, 10000 permutations).

**Fig. S2.**
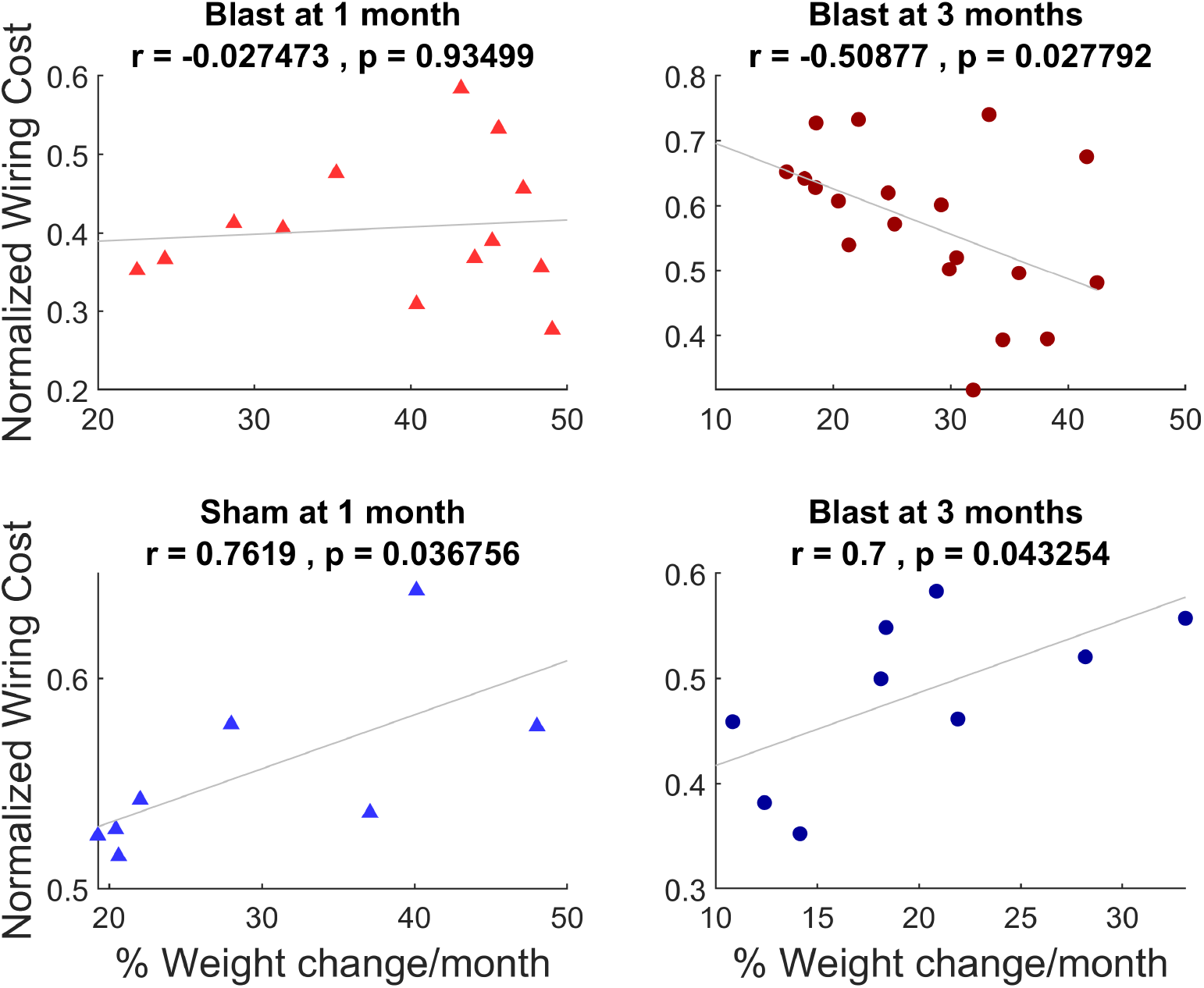
Opposite correlation signs between normalised network cost and normalised weight gain between injury groups in the chronic phase is robust when considering all four subgroups. **Top left:** blast at 1 month, **Top right:** blast at 3 months, **Bottom left:** sham at 1 month, **Bottom right:** sham at 3 months **All panels:** Spearman correlation values and associated *p*-values are reported. Lines are best fit lines using least-squares. (blast 1-month: n = 13, blast 3-month: n = 20, sham 1-month n= 8, sham 3 -month n = 9).

**Fig. S3.**
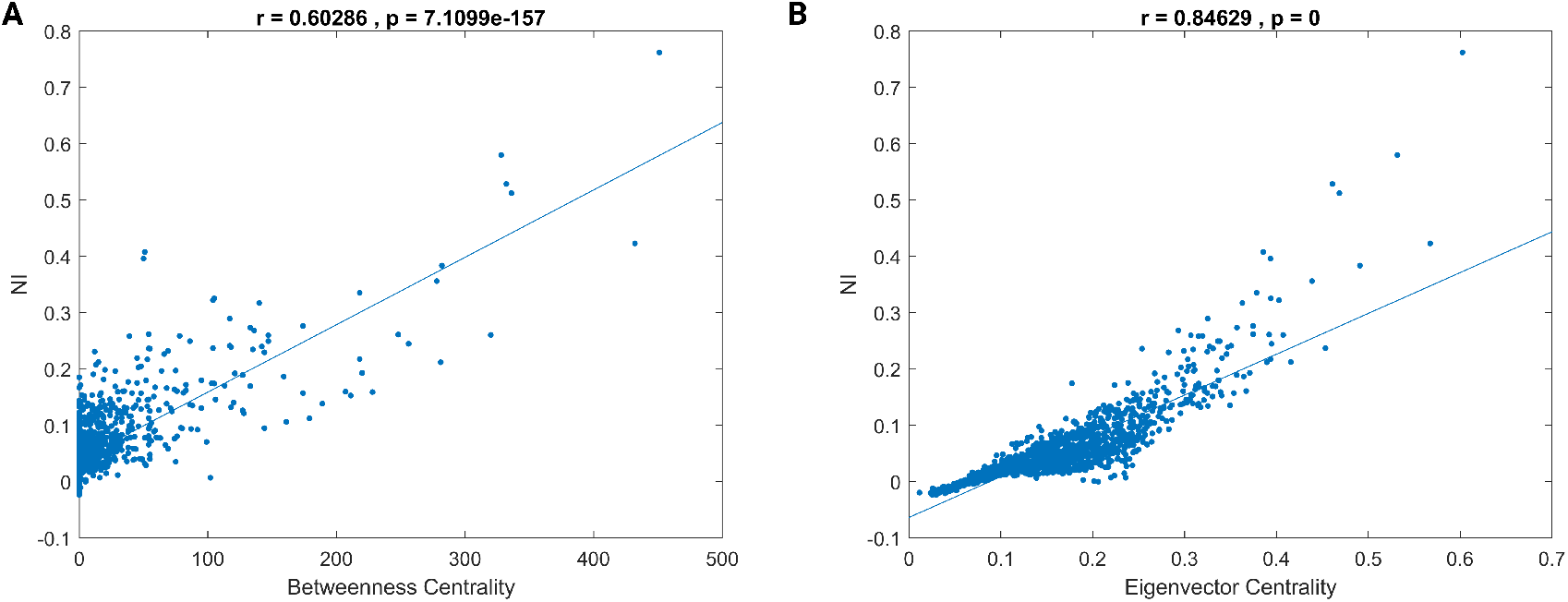
Node ictogenicity correlates with centrality measures at the node level **(A)** Betweenness centrality **(B)** Eigenvector centrality **All panels:** Spearman correlation values and associated *p*-values are reported.

**Fig. S4.**
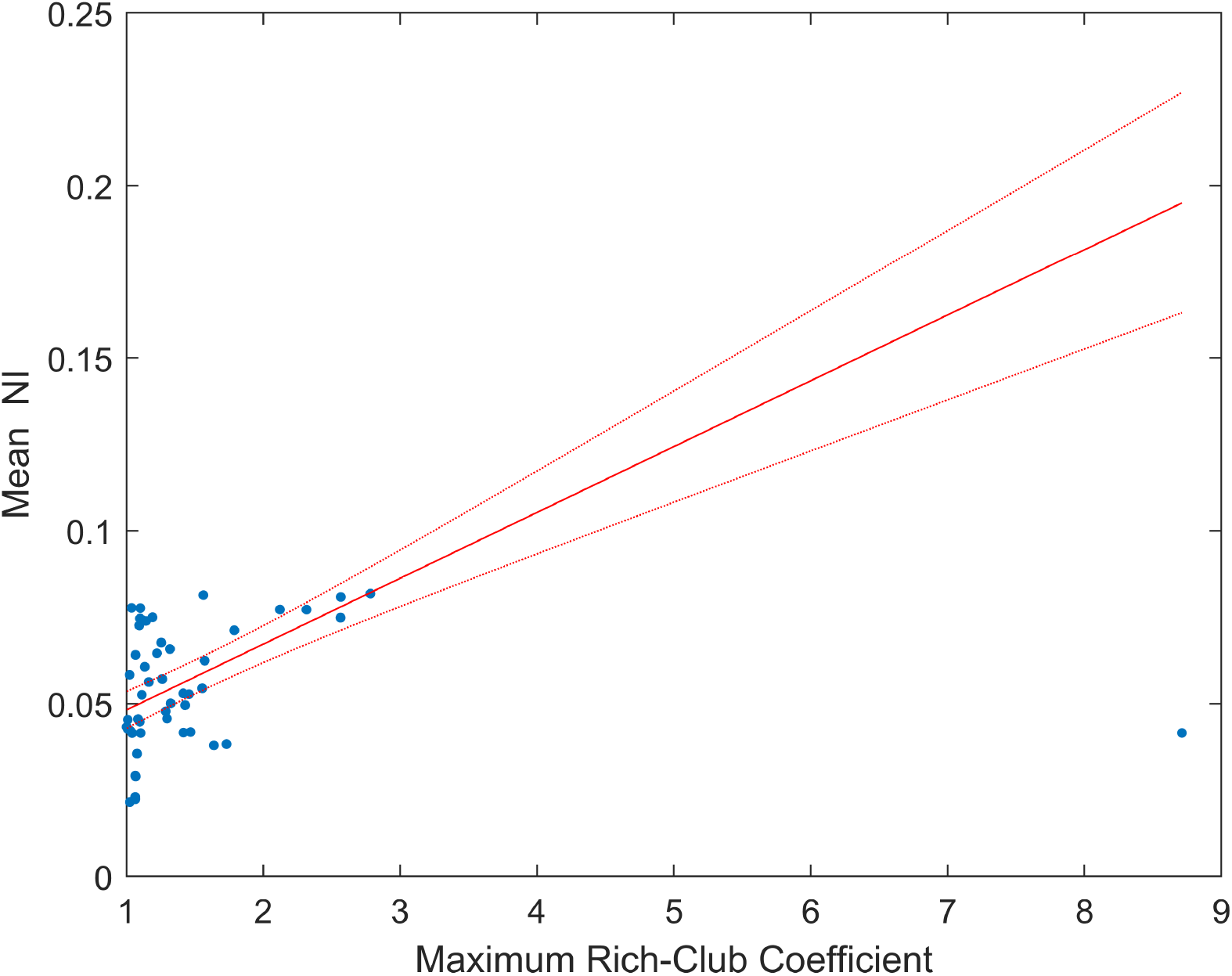
Mean Node ictogenicity correlates with rich-club presence at the network/subject level. Solid red line represents the line of best fit using robust regression, while dotted red lines represent the 95% confidence bounds.

**Fig. S5.**
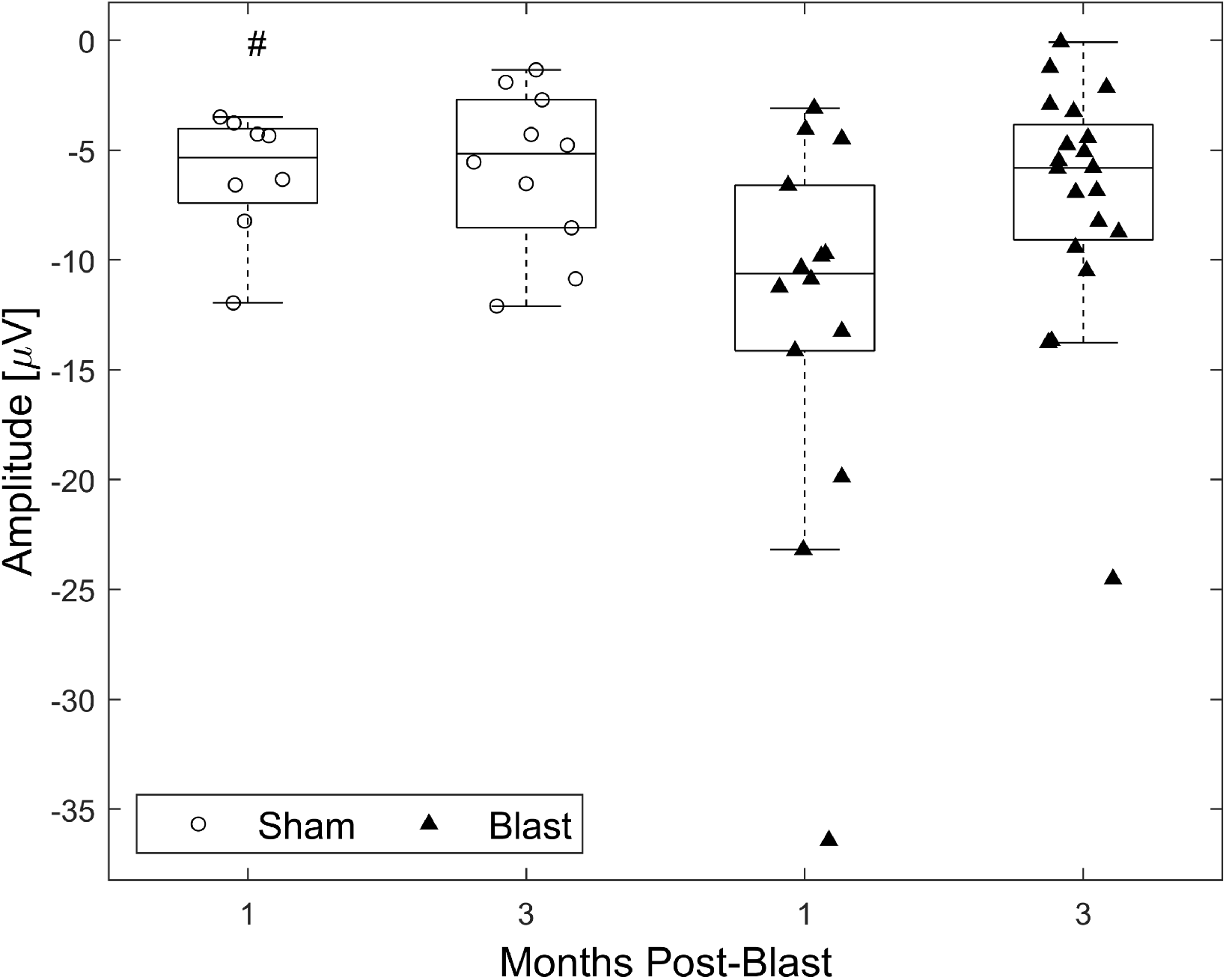
Increased AEP N1 amplitudes following blast in response to 10 kHz tone pips. Permutation-based 2-way ANOVA (Group x Time Post-Blast with interaction, 10000 permutations). Symbols denote *p*-values for a main effect of group. #: *p <* 0.05 (blast 1-month: n = 14, blast 3-month: n = 20, sham 1-month: n= 8, sham 3 -month: n = 10).

**Fig. S6.**
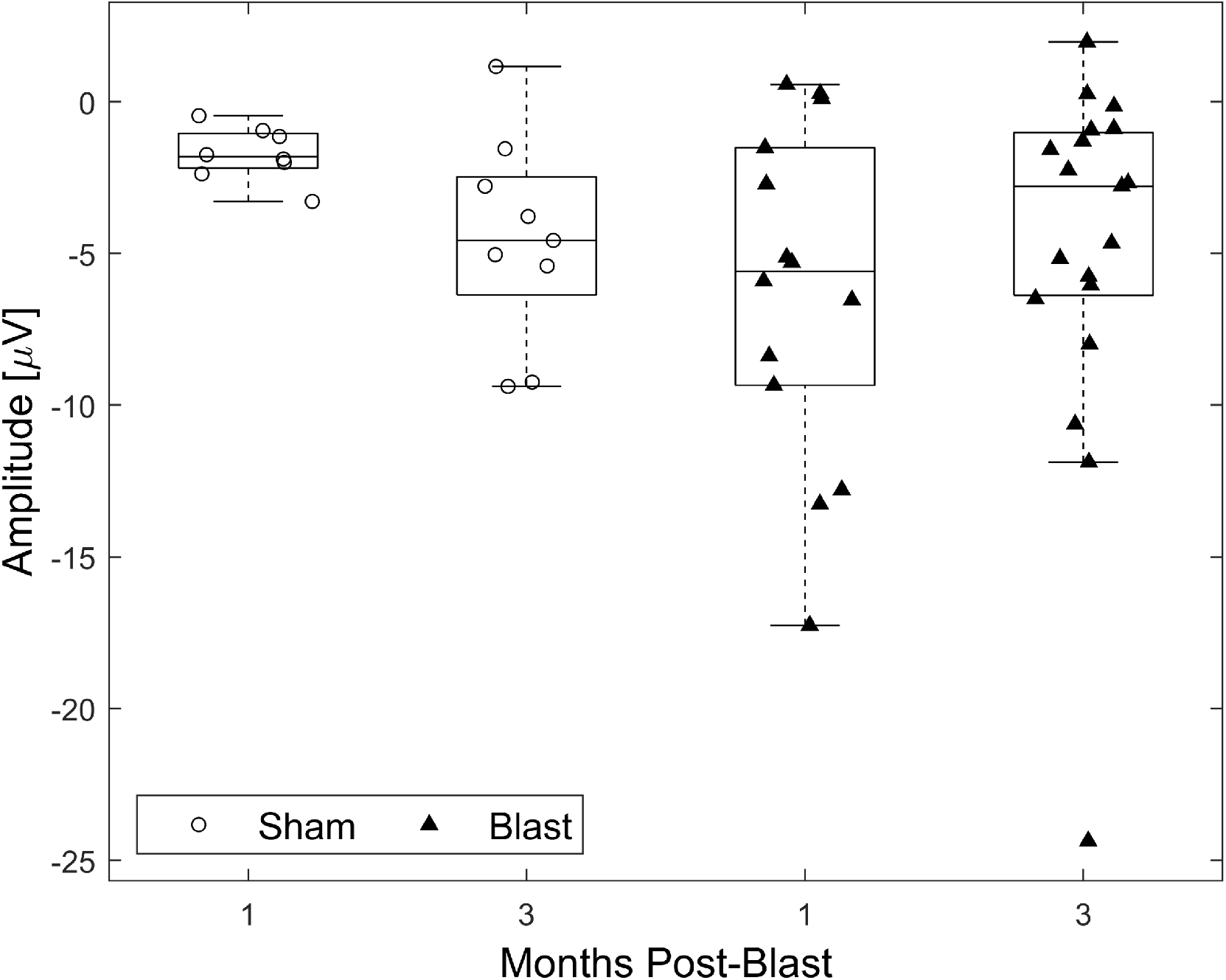
No effect of blast on AEP N1 amplitude in response to 37.5 kHz tone pips. Permutation-based 2-way ANOVA (Group x Time Post-Blast with interaction, 10000 permutations). (blast 1-month: n = 14, blast 3-month: n = 19, sham 1-month: n= 8, sham 3 -month: n = 9).

**Fig. S7.**
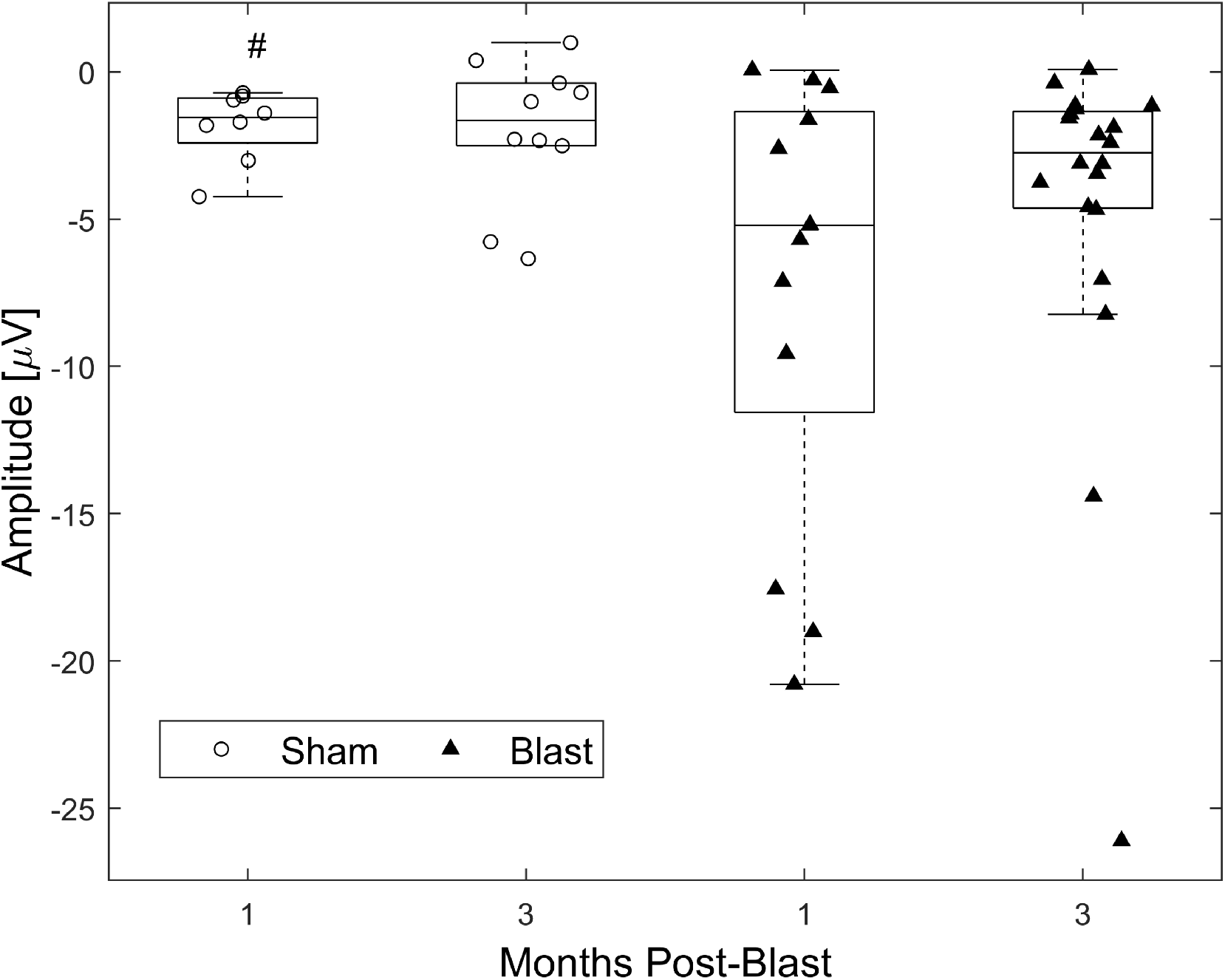
Increased AEP N1 amplitudes following blast in response to 65 kHz tone pips. Permutation-based 2-way ANOVA (Group x Time Post-Blast with interaction, 10000 permutations). Symbols denote *p*-values for a main effect of group. #: *p <* 0.05 (blast 1-month: n = 13, blast 3-month: n = 20, sham 1-month: n= 8, sham 3 -month: n = 10).

